# Accurate Variant Classification in Tumour-Only Genomic Data Using Interpretable Tabular Models

**DOI:** 10.64898/2025.12.09.693348

**Authors:** Lorenzo Tattini, Yiqing Yan, Nimisha Chaturvedi, Raja Appuswamy

## Abstract

Recent work has shown that machine learning can provide a reliable tool to classify somatic and rare germline variants in cancer studies where matched-normal samples are not available. Here, we present a workflow that combines an opensource pipeline with three machine-learning models, XGBoost, LightGBM, and TabNet, trained on eight types of features. Our approach substantially enhances the accuracy across all tested models providing accurate results irrespective of sample ancestry and tumour type. We build a parsimonious model and demonstrate that training on low-coverage data retains high accuracy when applied to high-coverage data and vice versa. In contrast to previous findings, our results indicate that XGBoost slightly outperforms LightGBM, achieving high classification accuracy even in the absence of copy-number information and allowing for the ancestry-unbiased calculation of the tumour mutational burden for different types of cancer.

## 1 Introduction

Accurate identification of somatic mutations is essential for cancer research. It is a critical aspect in clinical settings since it enables the characterisation of tumour biology and supports the application of biomarkers such as the tumour mutational burden (TMB) to guide immunotherapy [1]. Somatic variant calling is most accurate when matched-normal control samples are available to distinguish somatic mutations from germline variants [2]. However, control samples may be missing due to failed quality control or budget constraints, leading to reliance on tumour-only data. In such cases, misclassification of germline variants can inflate the TMB estimates, especially in individuals from under-represented populations in germline databases, thus introducing serious equity and reliability concerns [3].

Recent studies have demonstrated that machine learning can substantially improve tumour-only somatic variant classification by leveraging features derived from variant calls and external annotations extracted from publicly available databases [4]. Unfortunately, these methods involve computationally intense pipelines and rely on proprietary software, limiting their adoption and reproducibility.

To address this gap, we developed a fully open-source, lightweight, and reproducible framework for classifying mutations in tumour-only whole-exome sequencing (WES) data. Building on the same public data recently introduced [4], we implemented a streamlined workflow that performs “ensemble variant calling” (based on two opensource variant callers, Mutect2 and VarDict) [5] and derives a concise set of eight interpretable types of features to train three machine-learning classifiers (XGBoost,[6] LightGBM,[7] and TabNet[8]). Remarkably, our approach bypasses the calculation of copy-number variants and complex modelling steps yet improving classification accuracy [9]. Despite using a smaller set of features and lower computational overhead, our models achieve higher accuracy compared to previous studies [4]. Moreover, we show that models trained on high- or low-coverage data generalise effectively across different sequencing depths, and that ancestry-related biases in the TMB estimations are mitigated. Notably, XGBoost slightly outperforms the other models in most of test settings, including high-coverage data, underscoring the strengths of gradient boosting for this specific classification task. Our results highlight that an open-source pipeline that follows the best practices for cancer genomics can be paired with well-chosen features and robust machine learning models to deliver fast, accurate, and equitable tumour-only somatic variant calling across different cancer types.

## 2 Methods

### 2.1 Data acquisition

All the data processed in this study are summarised in Supplementary Table 1. The metastatic melanoma data were downloaded in FASTQ format from the “Sequence Read Archive” (https://www.ncbi.nlm.nih.gov/sra/docs/) using fasterq-dump 3.0.7 (“fasterq-dump –split-files $ind i”, where $ind i is the run ID, e.g. “SRR3083839”). All the other sequencing experiments were downloaded in BAM format from the TCGA data portal (https://portal.gdc.cancer.gov/) with the “gdc-client” command of the GDC data transfer tool. Each file was converted in two FASTQ files with “samtools fastq”. The human reference genome GRCh38 (https://ftp.ncbi.nlm.nih.gov) patch 14 was used for all the downstream analyses.

**Table 1.**
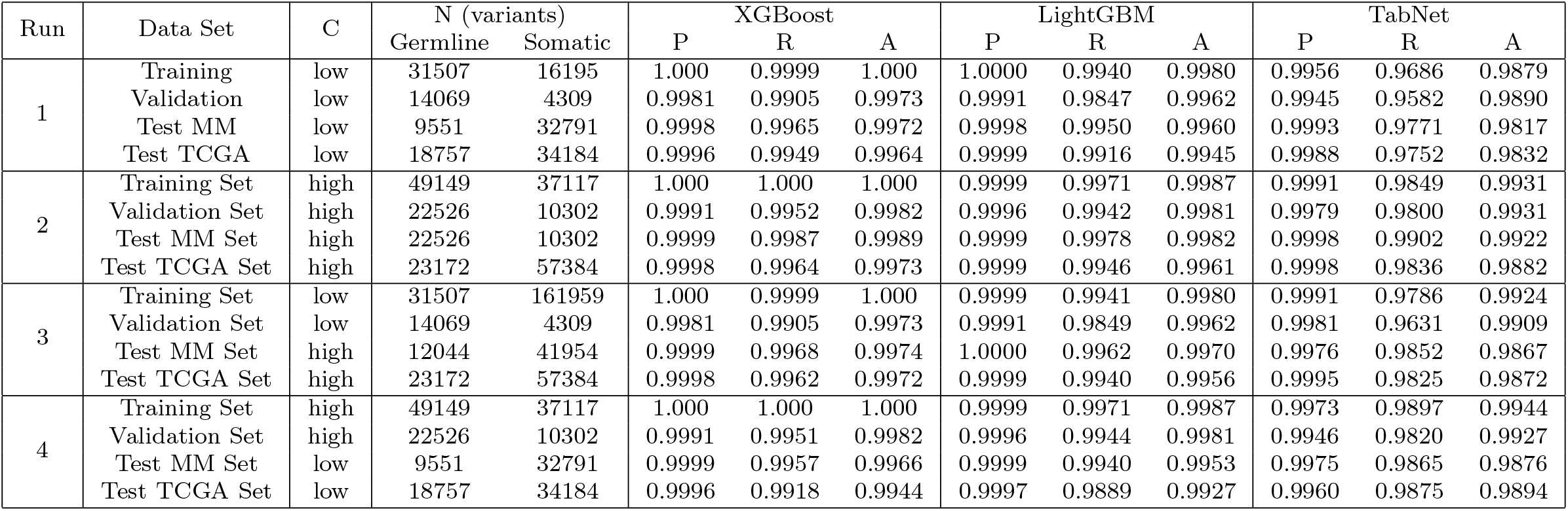
Precision (P), recall (R), and accuracy (A) of the three models run in different coverage (C) conditions on all the data sets. The numbers germline and somatic variants detected with the pipeline is also reported.

| Run | Data Set | C | N (variants) |  | XGBoost |  |  | LightGBM |  |  | TabNet |  |  |
| --- | --- | --- | --- | --- | --- | --- | --- | --- | --- | --- | --- | --- | --- |
|  |  |  | Germline | Somatic | P | R | A | P | R | A | P | R | A |
| 1 | Training | low | 31507 | 16195 | 1.000 | 0.9999 | 1.000 | 1.0000 | 0.9940 | 0.9980 | 0.9956 | 0.9686 | 0.9879 |
|  | Validation | low | 14069 | 4309 | 0.9981 | 0.9905 | 0.9973 | 0.9991 | 0.9847 | 0.9962 | 0.9945 | 0.9582 | 0.9890 |
|  | Test MM | low | 9551 | 32791 | 0.9998 | 0.9965 | 0.9972 | 0.9998 | 0.9950 | 0.9960 | 0.9993 | 0.9771 | 0.9817 |
|  | Test TCGA | low | 18757 | 34184 | 0.9996 | 0.9949 | 0.9964 | 0.9999 | 0.9916 | 0.9945 | 0.9988 | 0.9752 | 0.9832 |
| 2 | Training Set | high | 49149 | 37117 | 1.000 | 1.000 | 1.000 | 0.9999 | 0.9971 | 0.9987 | 0.9991 | 0.9849 | 0.9931 |
|  | Validation Set | high | 22526 | 10302 | 0.9991 | 0.9952 | 0.9982 | 0.9996 | 0.9942 | 0.9981 | 0.9979 | 0.9800 | 0.9931 |
|  | Test MM Set | high | 22526 | 10302 | 0.9999 | 0.9987 | 0.9989 | 0.9999 | 0.9978 | 0.9982 | 0.9998 | 0.9902 | 0.9922 |
|  | Test TCGA Set | high | 23172 | 57384 | 0.9998 | 0.9964 | 0.9973 | 0.9999 | 0.9946 | 0.9961 | 0.9998 | 0.9836 | 0.9882 |
| 3 | Training Set | low | 31507 | 161959 | 1.000 | 0.9999 | 1.000 | 0.9999 | 0.9941 | 0.9980 | 0.9991 | 0.9786 | 0.9924 |
|  | Validation Set | low | 14069 | 4309 | 0.9981 | 0.9905 | 0.9973 | 0.9991 | 0.9849 | 0.9962 | 0.9981 | 0.9631 | 0.9909 |
|  | Test MM Set | high | 12044 | 41954 | 0.9999 | 0.9968 | 0.9974 | 1.0000 | 0.9962 | 0.9970 | 0.9976 | 0.9852 | 0.9867 |
|  | Test TCGA Set | high | 23172 | 57384 | 0.9998 | 0.9962 | 0.9972 | 0.9999 | 0.9940 | 0.9956 | 0.9995 | 0.9825 | 0.9872 |
| 4 | Training Set | high | 49149 | 37117 | 1.000 | 1.000 | 1.000 | 0.9999 | 0.9971 | 0.9987 | 0.9973 | 0.9897 | 0.9944 |
|  | Validation Set | high | 22526 | 10302 | 0.9991 | 0.9951 | 0.9982 | 0.9996 | 0.9944 | 0.9981 | 0.9946 | 0.9820 | 0.9927 |
|  | Test MM Set | low | 9551 | 32791 | 0.9999 | 0.9957 | 0.9966 | 0.9999 | 0.9940 | 0.9953 | 0.9975 | 0.9865 | 0.9876 |
|  | Test TCGA Set | low | 18757 | 34184 | 0.9996 | 0.9918 | 0.9944 | 0.9997 | 0.9889 | 0.9927 | 0.9960 | 0.9875 | 0.9894 |

### 2.2 WES data processing and ensemble variant calling

The FASTQ files were processed with “mulo-wesml”, a publicly available pipeline (https://github.com/lt11/mulo-wesml). Briefly, mulo-wesml performed the following steps: (1) quality check with FastQC (version 0.12.1), (2) reference (hg38) indexing with “bwa index” (version 0.7.17), “samtools faidx” (version 1.18), and “gatk CreateSequenceDictionary” (version 4.5.0.0), (3) mapping with “bwa mem” [10], mate fixing with “samtools fixmate”, sorting with “samtools sort”, and duplicates marking with “samtools markdup”, and (4) coverage calculation with “samtools depth” [11]. The BAM files of each sample were used to call small variants, i.e. single-nucleotide variants (SNVs) and short insertions and deletions (indels) in matched mode with both (step 5a) VarDict (version 1.8.3) and (step 6a) GATK Mutect2 (version 4.5.0.0) using the panel of normals bundled within GATK. Variant calling was performed on target regions with a 100 bp padding, setting a minimal allele frequency threshold at 0.2 and a minimal depth at variant sites of 10 reads. Variants were then normalised (step 7a) with “vt normalize” (version 0.5) and de-duplicated with “vt uniq” [12]. The resulting variants were hard-filtered (step 8a) with “bcftools filter” (version 1.18) selecting only loci bearing at least one variant with TLOD ≥ 13 [11]. The TLOD value was automatically calculated within Mutect2. The intersection of the two sets (one set of variants from GATK and the other one from VarDict) was retained using “bcftools isec” with exact allele match. Moreover, only the variants annotated by VarDict as “LikelySomatic” and “StrongSomatic”, were retained. The VCF files were then sorted with “bcftools sort” and indexed with “tabix” (version 1.18) [13]. Finally, (step 9a) the variants were annotated in “cancer mode” with SnpEff and SnpSift (version 5.2.0) [14]. The latter was used to integrate in the VCF files the variant data available in dbNSFP (version 4.1.0) [15], cosmic (version 99), and dbSNP (version 155). The BAM files of tumour samples were also processed in unmatched mode with similar steps. SNVs and indels were called in unmatched mode with both (5b) VarDict and (6b) GATK Mutect2. Ensemble variant calling was performed on target regions with a 100 bp padding, setting a minimal allele frequency threshold at 0.2. The variants were then normalised with (7b) “vt normalize” (version 0.5) and de-duplicated with “vt uniq”. The resulting variants were hard-filtered (step 8b) with “bcftools filter” (version 1.18) selecting only loci bearing at least one variant with TLOD ≥ 13. The TLOD value (defined as the “log10 likelihood ratio score of variant existing versus not existing”) was automatically calculated within Mutect2. The intersection of the two sets (one set of variants from GATK and the other one from VarDict) was retained using “bcftools isec” with exact allele match. The VCF files were then sorted with “bcftools sort” and indexed with “tabix” (version 1.18). Finally, (step 9b) the variants were annotated with SnpEff and SnpSift (version 5.2.0). The latter was used to integrate in the VCF files the variant data available in dbNSFP (version 4.1.0), cosmic (version 99), and dbSNP (version 155). The gold standard VCF files were assembled by combining the variants called in matched mode (collected at step 9a) with those from the unmatched mode (collected at step 9b). Variants found in the former were labelled as “somatic”, while those reported in the latter were marked as “germline”. If the same variant was detected in both files only the call of the matched mode was retained.

### 2.3 Features engineering

We extracted eight types of features from the VCF files obtained from our pipeline. These included: (1) the number of reads at the variant site “DP”, (2) the fraction of reads supporting to the ALT allele “AF”, (3) the fraction of reads corresponding to the most abundant allele at variant loci “AD”, (4) the number of times an allele has been observed in the Catalogue Of Somatic Mutations In Cancer (COSMIC) “CNT”, (5) the maximum population frequency of the allele across multiple germline databases (all those reported in dbNSFP) “pop max”, (6) the ontology of mutations (missense, nonsense, disruptive inframe deletion, frameshift indel, non-coding) “ontology”, (7) the three-nucleotide context in the reference genome for single-nucleotide variants (1 bp upstream, the locus itself, and 1 bp downstream) “trinucleotide”, and (8) the type of transition or transversion single-nucleotide substitution “subs type”. To ensure that the workflow remained lightweight and easily deployable, we intentionally avoided incorporating features derived from copy-number variants. Overall, these eight types of features accounted for 89 binary features: DP, AF, AD, CNT, pop max, 6 ontologies, 65 trinucleotide contexts, and 13 substitution types.

### 2.4 BAM files downsampling

The final BAM files were downsampled with “samtools view -s $my frac -b $ind b”, where $my frac is the fraction of reads to be sampled to reach the desired mean coverage and $ind b is the input BAM file.

### 2.5 Model training

We implemented a Python (version 3.10.12) workflow to extract variant-level features from the annotated single-sample VCF files and train supervised learning models for the classification of somatic and germline variants. The VCF files were parsed using pandas (version 2.2.3), and variant annotations (described in section 2.3) were extracted from the INFO field except the AD annotation that was extracted from each sample column. Functional categories were assigned through a predefined ontology mapping encompassing missense, nonsense, in-frame indel, frameshift indel, and non-relevant (NR) variants (e.g. intronic variant, intergenic variants). Population allele frequencies were derived by parsing dbNSFP entries (e.g., ExAC, 1000Gp3, ESP6500) to compute a maximum population frequency (pop max), which was subsequently used to filter out common germline variants (pop max ≥ 0.01). Additional filtering excluded germline variants that were not annotated with relevant ontology labels. The latter included:

- stop_lost
- start_lost
- missense_variant
- missense_variant&splice_region_variant
- stop gained
- frameshift variant&stop_gained
- stop_gained&splice_region_variant
- disruptive inframe deletion
- conservative_inframe deletion
- conservative_inframe insertion
- frameshift_variant&splice_donor variant&splice_region_variant&intron_variant
- frameshift_variant&splice_region_variant
- frameshift_variant&stop_lost&splice_region_variant
- frameshift_variant

Feature vectors included numeric predictors (DP, AF, AD, CNT, pop max, nb variants) and categorical ones (subs type, trinucleotide, ontology), which were one-hot encoded using the category encoders library.

The resulting matrices were partitioned into training, validation, and test sets, each derived from distinct tumour subtypes (as discussed in section 3.1). Three machine-learning algorithms were benchmarked using identical input features. XGBoost (version 2.1.4) was run with parameters learning rate=0.05, max depth=30, n estimators=750, objective: binary:logistic) wihle LightGBM (version 4.6.0) with num leaves=31, learning rate=0.05, feature fraction=0.9, boosting type=gbdt, objective: binary). The TabNet architecture (from pytorch-tabnet version 4.1.0) was defined with parameters n d=24, n a=24, n steps=4, lamba sparse=10^-4^, momentum=0.3, clip value=2.0, optimizer fn=torch.optim.Adam, scheduler params= }gamma:0.95, step size:20}, scheduler fn=torch.optim.lr scheduler.StepLR, epsilon=10^-15^). The TabNet classifier was trained with default learning rate (0.02) for up to 100 epochs (max epochs=100) with an early stopping patience of 100 epochs (patience=100), using a mini-batch size of 4000 and a virtual batch size of 256. All the models were trained on CPU in less than 2 hours in total.

Model performance was evaluated using the area under the receiver operating characteristic (ROC) curve, computed via scikit-learn (version 1.6.1). Feature importance was estimated directly from model gains (XGBoost and LightGBM) or attention masks (TabNet) and visualized using matplotlib (version 3.8.4) and seaborn (version 0.13.2).

### 2.6 Variant classification and performance evaluation

We calculated precision, TP / (TP + FP), recall TP / (TP + FN), true negative rate (TNR), TN / (TN + FP), negative predictive value (NPV), TN / (TN + FN), and accuracy (TP + TN) / (TP + TN + FP + FN) by defining a gold standard based on our combined (matched-normal and tumour-only) pipeline: somatic variants identified in the matched-normal analysis were considered true positives if correctly predicted by the model and false negatives if missed, while germline variants were labelled as true negatives if properly classified and false positives if erroneously predicted as somatic. This evaluation framework ensured that performance metrics reflected the capability of a model to distinguish genuine somatic mutations from rare germline variants, thereby providing a robust measure of classification accuracy and reliability.

### 2.7 Statistical analyses

Samples that did not provide any variant in the matched pipeline due to sequencing issues were filtered out (fastq file IDs: 18382739a33f49e6856ffcf720c49363, 693ee45f0cc945e18b05bfb08253990a, and fecf5edc7a044f9da47820c262780124). All the statistical analyses were performed in R (version 4.3.2) using Rstudio (024.12.0+467).

### 2.8 Tumour mutational burden calculation

The TMB was calculated counting all the relevant variants in the SnpEff annotations. These included:

- stop_lost
- start _lost
- missense _variant
- missense _variant&splice _region_ variant
- stop _gained
- frameshift_ variant&stop _gained
- stop _gained&splice _region_ variant
- disruptive _inframe _deletion
- conservative _inframe _deletion
- conservative inframe _insertion
- frameshift_ variant&splice _donor _variant&splice _region_ variant&intron_ variant
- frameshift_ variant&splice _region_ variant
- frameshift_ variant&stop _lost&splicev _region_ variant
- frameshift_ variant

The count was normalised by the size in Mbp of the regions targetted in the corresponding WES kit. The kits used were: HGSC VCRome (45.1 Mbp), NimbleGen SeqCap EZ Exome V3 (151.7 Mbp), Agilent Custom V2 (56.8 Mbp).

## 3 Results

### 3.1 Overview of data sets, pipelines, features, and models

In order to boost the accuracy of the prediction of somatic variants from tumour-only variant calling, we implemented a mutation classification pipeline that processes samples with and without matched-normal sequences and leverages these data to train three machine learning models.

We curated a diverse set of 215 tumour samples (Supplementary Table 1), and their matched-normal samples, covering a broad spectrum of biological characteristics already used for this type of application [4]. These included high-purity, moderately low-TMB ovarian adenocarcinoma, low-purity and low-TMB stomach adenocarcinoma, as well as metastatic melanoma, and lung adenocarcinoma and squamous carcinoma, both known for high TMB. We organised the data in a training set comprising 103 tumour samples from: 15 bladder urothelial carcinoma (BLCA) [16], 13 glioblastoma multiforme (GBM) [17], 15 head and neck squamous cell carcinoma (HNSC) [18], 15 lung adenocarcinoma (LUAD) [19], 15 lung squamous cell carcinoma (LUSC) [20], 15 stomach adenocarcinoma (STAD) [21], and 15 ovarian serous cystadenocarcinoma (OV) [22]. The validation set consisted of 15 samples from colon adenocarcinoma (COAD) [23], 15 lymphoid neoplasm diffuse large B-cell lymphoma (DLBC), and 14 testicular germ cell tumours (TGCT) [24]. The two test sets included: (1) 23 metastatic melanoma (MM) tumour samples [25] and (2) a TCGA set that comprised 15 breast invasive carcinoma (BRCA) [26], 15 sarcoma (SARC) [27], 15 uterine corpus endometrial carcinoma (UCEC) [28]. The selection of these subtypes aimed to capture the variability In mutation rates, copy-number landscapes, and purity levels, providing a comprehensive framework for training, validating, and testing robust machine-learning models. All patient-specific metadata, including cancer subtype, TMB, purity, CNV burden, and ancestry, are reported in Supplementary Table 1 and were retrieved from McLaughlin *et al*. [4].

We designed eight mutation-specific types of features that could be retrieved in tumour-only samples (Figure 1). This set of features was designed to efficiently distinguish between somatic and germline variants, optimising performance for tumour-only variant calling. The selected features included traditional indicators for somatic variant detection, such as germline database frequency, COSMIC somatic mutation database counts, and read-based statistics like variant allele fraction and major allele frequency. Additionally, we integrated features representing the trinucleotide context and base substitution subtypes to capture mutational signature patterns that may distinguishing somatic mutations from germline variants.

**Fig. 1.**
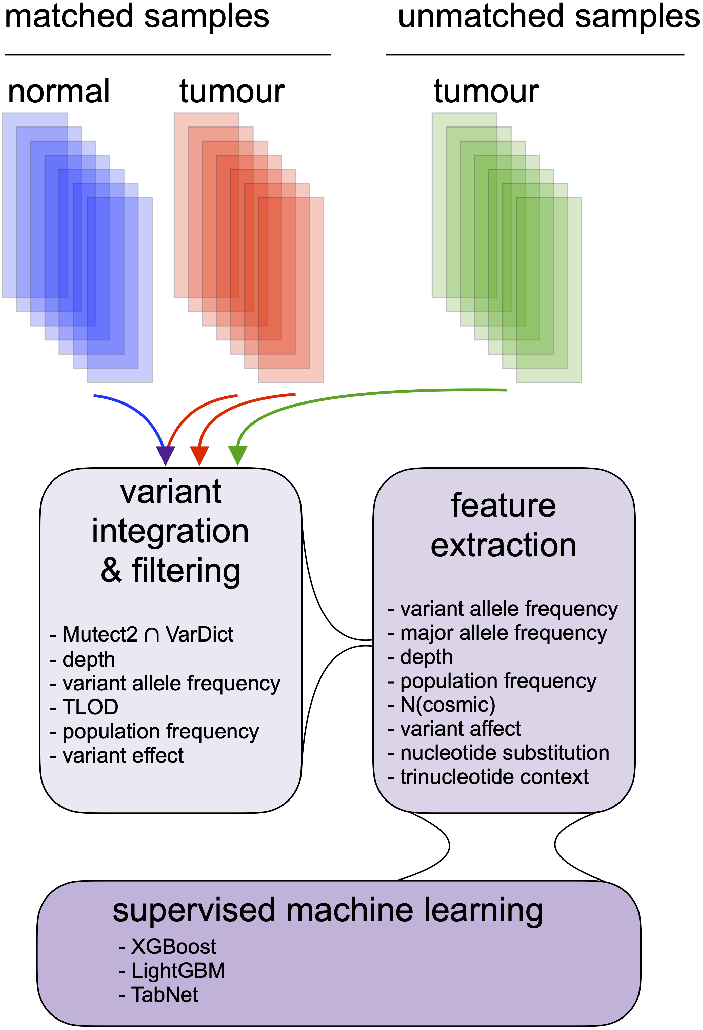
Overview of the workflow. We developed a framework that can process both matchednormal and tumour-only data. When running the matched-normal pipeline we call variants both in matched-normal mode and in tumour-only mode. The two sets are then integrated to build the gold standard, retaining: (1) among duplicates only the variants called in matched-normal mode (with their somatic labels) plus (2) all the other variants called in tumour-only mode. The latter are labelled as germline. Variants are filtered to remove common variation observed in previous studies and reported in germline databases. Eight types of features are then extracted and used to train three models (XGBoost, LightGBM, and TabNet) via supervised machine learning.

The somatic truth labels were determined through an open-source variant-calling pipeline using matched-normal samples. Remarkably, our pipeline exploits only opensource software, and is freely available (https://github.com/lt11/mulo-wesml). The ensemble variant calling pipeline includes two variant callers, Mutect2 and VarDict that can operate both in matched-normal and in tumour-only mode [29]. Variants identified as somatic in the matched-normal pipeline were labelled accordingly, and merged with variants obtained from tumour-only data. The latter were labelled as germline, as previously reported [4]. When duplicate variants were detected only the variant called in matched-normal mode was retained. This binary classification framework formed the foundation for training our supervised machine learning models (Figure 1).

We employed three machine-learning models tailored for tabular data: TabNet,[8] an attentive deep-learning model, and two gradient boosting methods: XGBoost,[6] a tree-based algorithm, and LightGBM,[7] a similar method optimized for speed and efficiency. Our approach focused on maximising predictive accuracy while reducing computational costs by leveraging only eight carefully selected types of features to achieve reliable somatic variant classification.

### 3.2 Building the full model from a toy model

As an initial exploration, we constructed a baseline “toy” model that incorporated only those features reported in the full model as having minimal or low predictive importance in tumour-only variant classification tasks. This baseline served both as a lower-bound reference and as a controlled starting point for systematically assessing the incremental contribution of each additional feature class. We then adopted a stepwise feature-addition strategy, progressively reintroducing higher-importance features one at a time according to their ranked relevance (Supplementary Table S2). We progressively added: subs type C*>*A, trinucleotide GTG, trinucleotide TGA, trinucleotide GGA, trinucleotide TCC, subs type G*>*C, ontology missense, subs type C*>*T, subs type G*>*T, trinucleotide GGG, trinucleotide TCA, subs type G*>*A, ontology nonsense, subs type NA, pop max, AD, ontology frame shift indel, AF, ontology inframe indel, DP, ontology NR, CNT.

As shown in Figure 2 at each step, we trained the model and quantified performance using the area under the ROC curve (AUC) and accuracy, computed independently for all validation and both test data sets. This design allowed us to characterise not only the overall discriminative power of the model, but also how rapidly performance converged toward an optimum as the feature set expanded. Across all data sets and models, we observed that AUC increased sharply during the first stages of feature addition and began to stabilise once 85 out of the 89 total features were included, indicating that the majority of discriminative signal is captured by this subset. In contrast, accuracy continued to exhibit small but measurable improvements up to the full 89-feature configuration. Nonetheless, after 85 features, accuracy already exceeded 0.95 for all data sets with one notable exception: the metastatic melanoma test set analysed with TabNet, for which the final incremental features contributed a modest additional gain. Although model performance exhibited considerable variability across data sets, both in terms of AUC and accuracy, when fewer than 85 features were used, this variability markedly decreased as the feature set grew. Once 85 or more features were included, the performance metrics consistently converged toward their respective optima for all data sets, indicating that the final subset of features provides the stability and discriminative power necessary for robust generalisation across tumour types and test conditions.

**Fig. 2.**
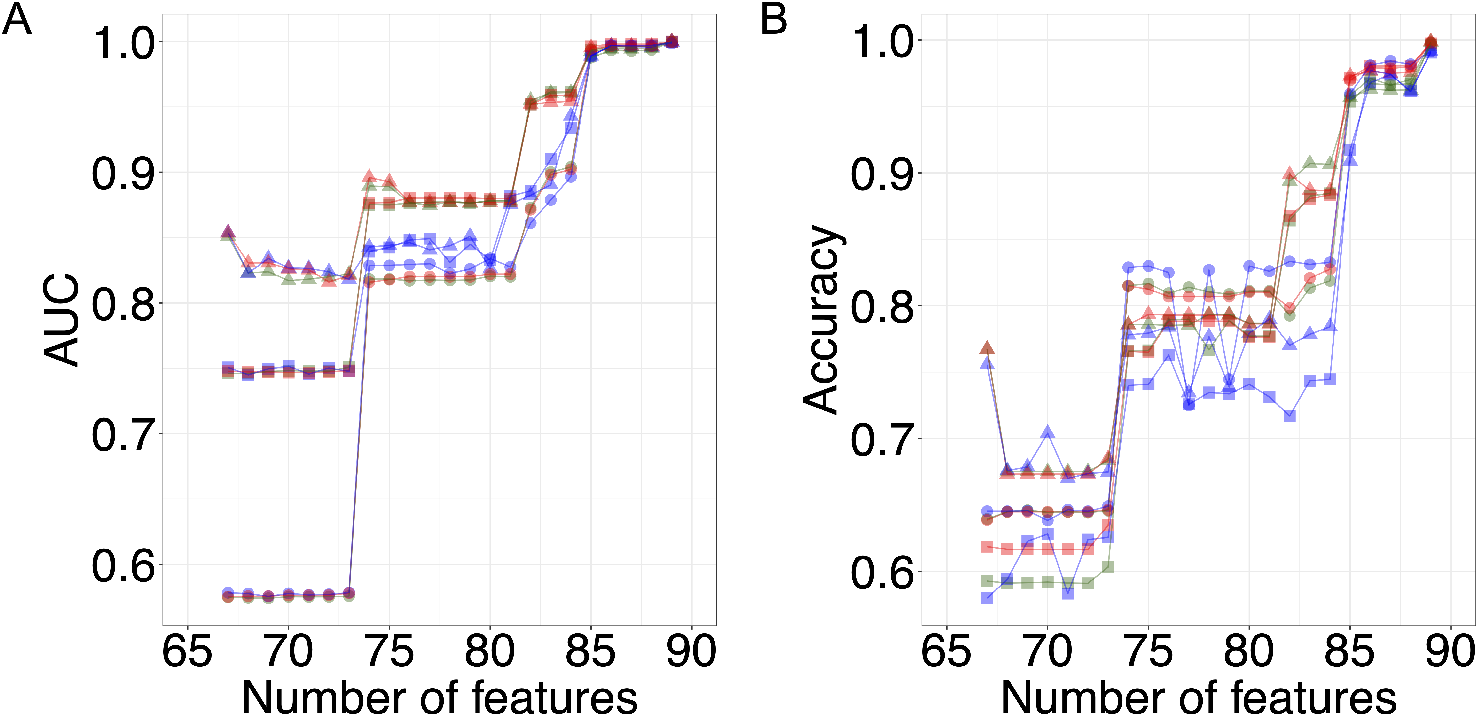
Area under the ROC (receiver-operating characteristic) curve (AUC) (A) and accuracy (B) as a function of the total number of features used to train the models: XGBoost (red), LightGBM (green), and TabNet (blue). The values for validation (circles), metastatic melanoma test (triangles), and TCGA test (squares) data sets are reported. Solid lines serve as an eye guide.

Taken together, these results indicate that while a reduced feature set of 85 features captures most of the predictive signal, the full 89-feature model systematically achieves the highest and most stable performance across metrics and data sets. For this reason, we selected the complete 89-feature configuration as the optimal model and employed it consistently in all subsequent analyses and evaluations.

### 3.3 Model accuracy with coverage-unbalanced sets and by type of variant

We calculated the confusion matrix considering somatic variants from the matched pipeline as positive elements and germline variants as negative elements. First, we evaluated the accuracy of the three models using high coverage data. All the three models achieved high performance with both test sets, showing accuracy larger than

0.99 and reflecting high precision, recall (Table 1, run 2), as well as high true negative rate (TNR), and negative predicted value (NPV) (Supplementary Table S3). The only exception was the TabNet which showed an accuracy of 0.98 with the TCGA test set due to a slight decrease in the recall value. On the other hand, precision was still very high (P *>* 0.99). Thus, in optimal conditions (high coverage training data and high coverage test data, Table 1 run 2) XGBoost provides a narrowly better performance over LightGBM and TabNet. We also tested our approach on low-coverage data (Table 1, run 1), the latter obtained by downsampling the high-coverage mapping files. Remarkably, also in this case all the three models provided very accurate results (A *>* 0.99) with both test sets. Again, the only exception was TabNet with A = 0.9817 for the MM test data set and A = 0.9832 for the TCGA test data set.

Then, we explored the robustness of our method by training with high-coverage data and testing with low-coverage samples (run 4) and vice versa (run 3). Both tree-based models achieved very high accuracy (*>* 0.99) while TabNet scored 0.9832 and 0.9802 with the MM data set and the TCGA data set, respectively. In order to further stress the models, we trained them with low-coverage data and tested on high-coverage samples. Strikingly, also in this case the performance was very high, achieving accuracy larger than 0.99 with both the tree-based models irrespective of the test data set used, while TabNet showed A = 0.9876 and A = 0.9894 with the MM data set and the TCGA data set, respectively. Using the MM test set, we observed a slight Increase of XGBoost accuracy when testing on high coverage data after training on low coverage data (A = 0.9974, run 3) compared to the training on high coverage data and testing on low coverage data (0.9966, run 4). An opposite effect was observed with LightGBM (A = 0.9970 and A = 0.9953, low coverage training for high coverage testing and vice versa, respectively, as shown in run 3 and run 4). Thus, tree-based models proved to be remarkably robust to different coverage values in the training and test sets. Also TabNet demonstrated strong robustness to differences in coverage between training and test sets, despite exhibiting a lower baseline accuracy, as discussed above. Moreover, we observed high accuracy irrespective of the type of variant analysed, SNVs or indels, as in both cases the accuracy observed was larger than 0.99 for all the metrics (Supplementary Table S4). The only exception was the precision and recall values observed for the indels of the metastatic melanoma test set (although with P 0.93, R, 0.95). The same trend was confirmed across all the models, suggesting that it was related to statistical fluctuations due to the low number of indels present (82).

Overall, we concluded that the tree-based gradient boost models implemented in our workflow provide slightly better performance compared to TabNet and that although both XGBoost and LightGBM can cope with coverage differences between the training and the test sets (Table 1, run 3 and 4), XGBoost provides the best performance in optimal conditions (Table 1, run 2). These results differ from previous reports [4] and highlight the impact of pipeline construction variant labelling.

### 3.4 Analysis of feature importance

Using high-coverage data, we investigated the importance of the features used to train the models (Figure 3). The “CNT” feature, representing the number of times a given allele has been reported in the Catalogue of Somatic Mutations in Cancer (COSMIC), emerged as the most influential predictor across all the models analysed. Additional features with notable contributions included “DP” (read depth at the variant locus), “ontology NR” (“o NR” for short, capturing low-impact functional categories such as synonymous, intronic, or intergenic variants), and indel-related ontology terms. Conversely, features encoding the trinucleotide sequence context and the substitution type exhibited consistently low importance values, irrespective of the model considered.

**Fig. 3.**
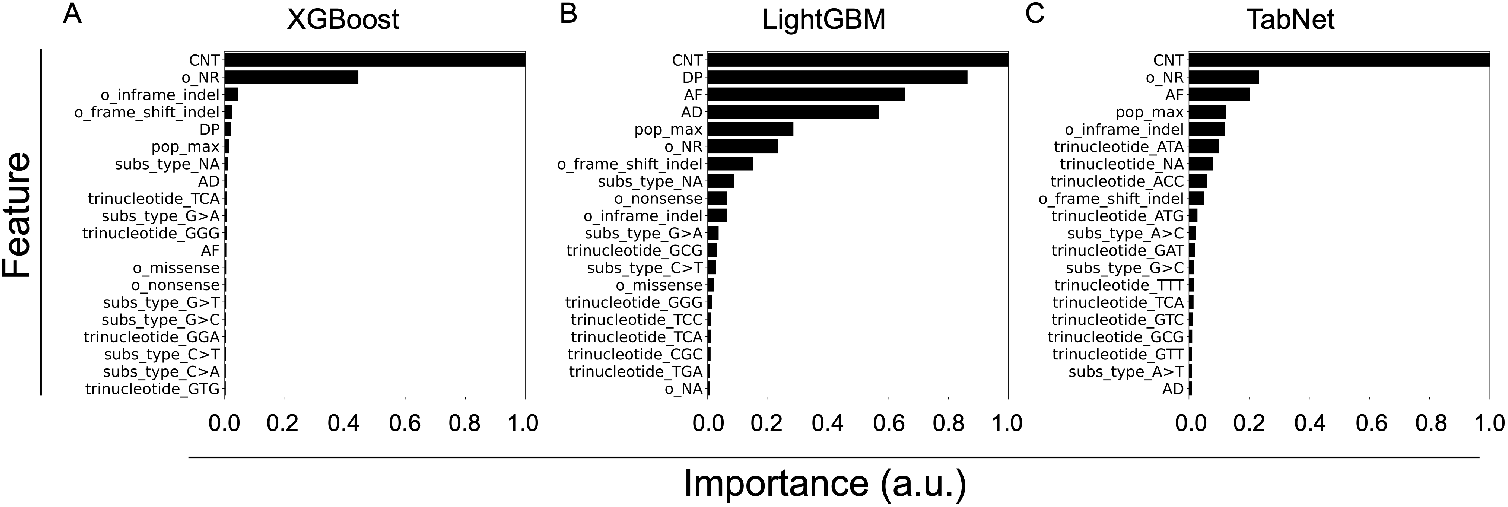
Relative importance of features in tabular models for somatic and rare germline variant classification. The 20 most important features are shown for (A) XGBoost, (B) LightGBM, and (C) TabNet. All the values were calculate from high-coverage data. Ontology-related features, e.g. “ontology NR”, are abbreviated as e.g. “o NR”.

Interestingly, XGBoost concentrated most of the importance on the top two features, while LightGBM and TabNet distributed feature importance more evenly across a broader set of predictors. Previous studies [4] had noted the unexpectedly low importance assigned to ontology-related features, which was surprising given the assumption that nonsense mutations are more likely to occur as somatic rather than germline events. Our models implementation seems to have partially solved this discrepancy, showing “missense” and “non-sense” features ranked in the top-20 although of tree-based models with low importance.

Finally, we assessed the robustness of the models to feature removal. We conducted a series of tests where key features were progressively removed, and the models were retrained and evaluated on high-coverage data. When the “CNT” feature was excluded, the average accuracy (across both the MM and TNBC test sets) decreased to approximately ∼ 0.98, ∼ 0.98, and ∼ 0.86 for XGBoost, LightGBM, and TabNet, respectively. The exclusion of the top four features led to a more pronounced drop in performance, with mean accuracies reaching ∼ 0.88, ∼ 0.90, and ∼ 0.50 for XGBoost, LightGBM, and TabNet, respectively. As expected, removing low importance features did not impact the accuracy of any of the models, leaving accuracy values on average unchanged (Supplementary Table S3).

These results indicate that both tree-based approaches exhibit strong robustness. Yet, LightGBM is slightly more resilient to feature removal than XGBoost, in agreement with their respective feature importance profiles. In contrast, TabNet is heavily affected in particular due to a dramatic decline of recall (on average from *∼* 0.98 to *∼* 0.66).

### 3.5 Tumour mutational burden

Accurate TMB estimation is a key aspect for any model aimed at improving tumour-only variant calling. This is particularly important in immuno-oncology clinical trials where TMB serves as a biomarker of treatment response and patient survival [30, 31].

Calculating the tumour mutational burden from low-coverage whole-exome sequencing (WES) data can result in systematic underestimation, primarily due to reduced sensitivity in variant detection. At lower sequencing depths, many true somatic mutations, especially those present at low variant allele fractions or in regions with poor coverage, may be missed by small variant callers. This leads to an incomplete representation of the tumour mutational landscape and may compromise the reliability of TMB as a predictive biomarker in clinical settings. The analysis of metastatic melanoma samples revealed that downsampling the BAM files from a mean coverage of 87 to 25 resulted in the loss of 50% of missense somatic variants, and 59% of frameshift somatic variants. Thus, in the following analyses we focused on highcoverage data only. First, we evaluated the Kendall rank correlation coefficients using the TMB obtained from the matched-normal pipeline and the TMB values from the somatic variants predicted by the models (Table 2). As expected, the two tree-based gradient boost models showed higher correlation with the gold standard data.

**Table 2.** Kendall rank correlation coefficient for the TMB calculated with the matched pipeline and the three models.

| Set | XGBoost | LightGBM | TabNet |
| --- | --- | --- | --- |
| MM | 1.0000 | 0.9913 | 0.9740 |
| TCGA | 0.9932 | 0.9871 | 0.9578 |

To further characterise the performance of our framework, we assessed whether TMB estimates remain robust across different tumour types. For each cancer subtype, we compared the TMB predicted by the XGBoost model with the TMB calculated using the matched tumour–normal pipeline (Supplementary Table S1). The resulting linear regressions (Figure 4) consistently showed strong concordance between predicted and matched-normal TMB values, indicating that our approach maintains reliable performance irrespective of tumour type.

**Fig. 4.**
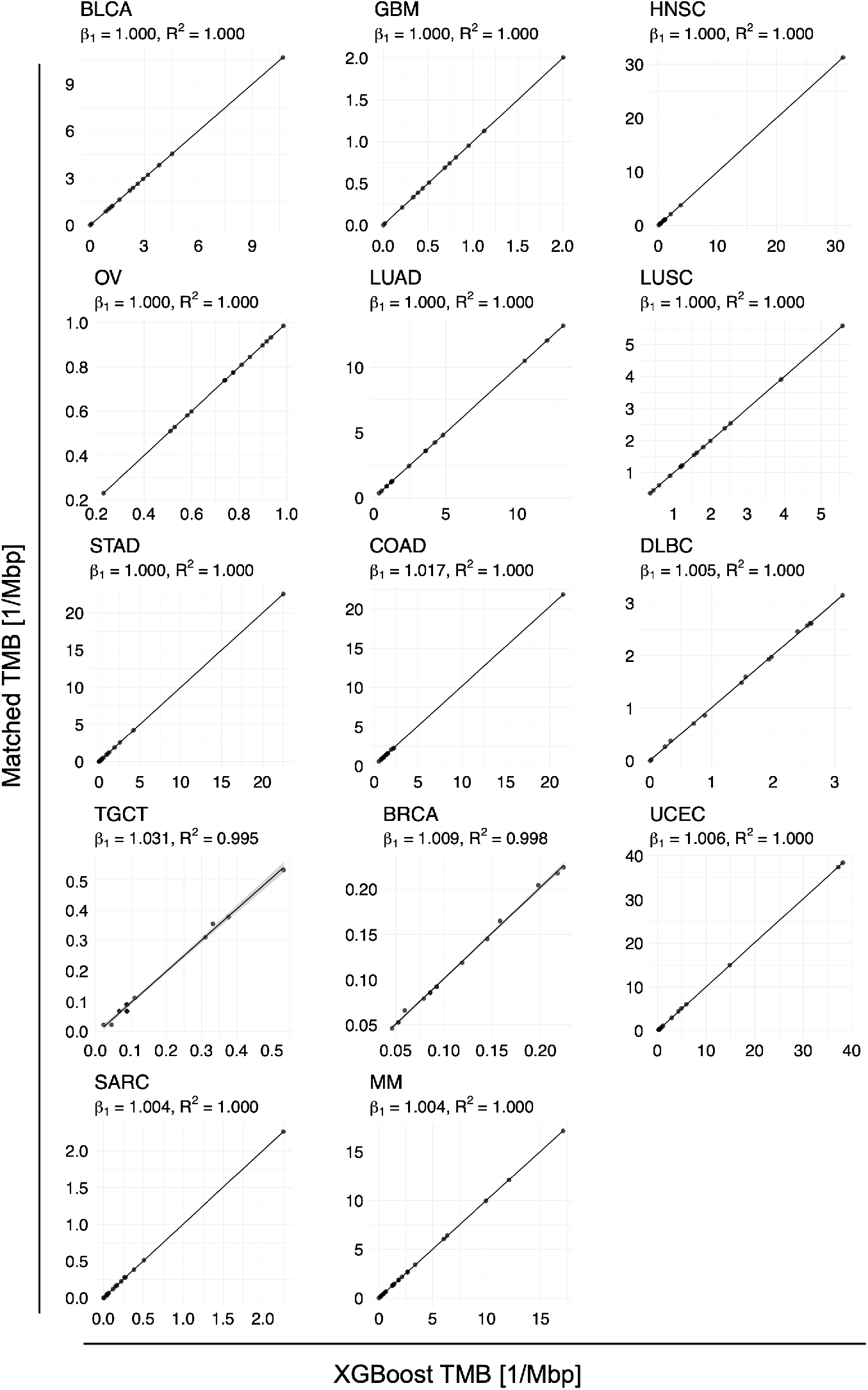
Linear regressions comparing the TMB calculated using the matched tumour–normal pipeline with the TMB predicted by the XGBoost model. Notably, the axes ranges differ among tumour types. The plots show the regressions for all the tumour type of the training set (bladder urothelial carcinoma (BLCA), glioblastoma multiforme (GBM), head and neck squamous cell carcinoma (HNSC), lung adenocarcinoma (LUAD), lung squamous cell carcino1m9a (LUSC), stomach adenocarcinoma (STAD), and ovarian serous cystadenocarcinoma (OV)), the validation set (colon adenocarcinoma (COAD), lymphoid neoplasm diffuse large B-cell lymphoma (DLBC), and testicular germ cell tumours (TGCT)), the TCGA test set (breast invasive carcinoma (BRCA), arcoma (SARC), uterine corpus endometrial carcinoma (UCEC)), and the MM test set.

### 3.6 Ancestry bias

Tumour-only variant calling is particularly vulnerable to disparities in germline databases, with recent studies showing that the absence of matched-normal samples leads to the inflation of TMB estimates in individuals from under-represented populations [3]. This under-representation of specific ancestries in genomic resources presents a critical barrier to equitable precision oncology as it may lead to biased biomarker detection and influence clinical decisions. Notably, machine-learning approaches that integrate multiple orthogonal features have shown promise in correcting these biases, enabling more accurate and equitable TMB estimation for patients from diverse populations.

For each of the ancestries in the test data sets, we calculated the TMB using the variants from the matched pipeline as well as the three models (Figure 5) and applied, for each ancestry, the Mann–Whitney U test to the TMB from the matched-normal data and each model. All the tests showed p-values larger than 0.95, thus failing to detect any significant difference between the TMB values from the matched pipeline and the TMB estimation from the models, and suggesting they may come from the same distribution.

**Fig. 5.**
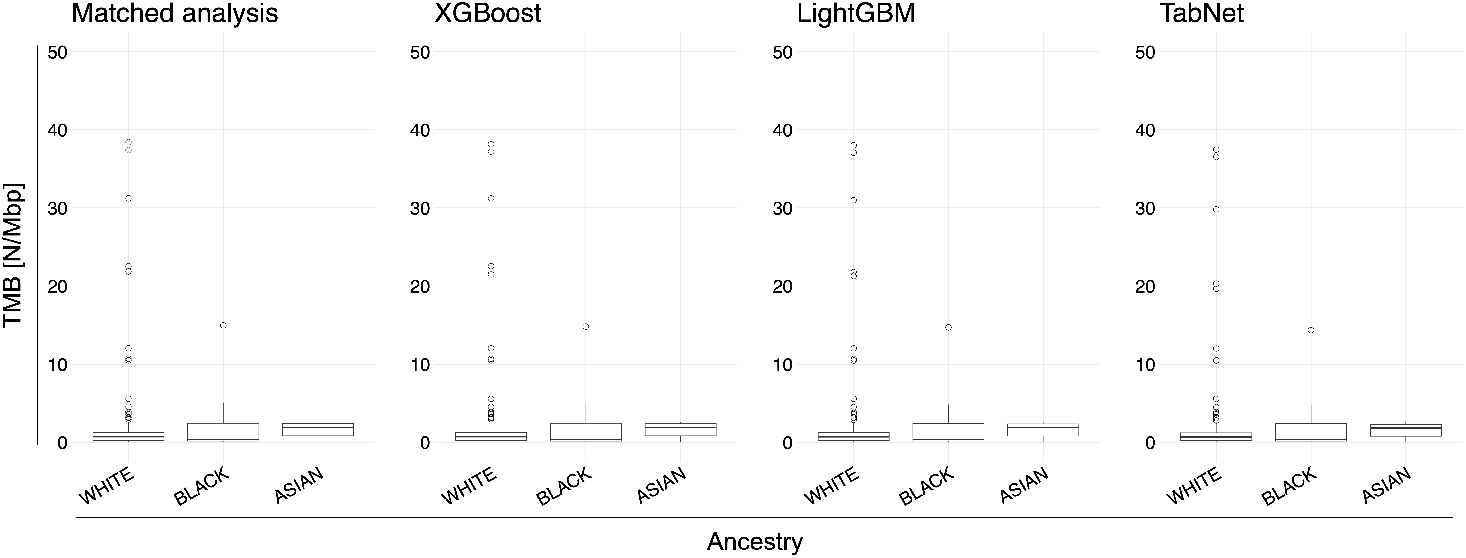
Tabular machine learning methods mitigate the ancestry-related bias in germline databases for TMB estimation in tumour-only WES analyses. The box plots of the TMB were calculated from: the matched pipeline, and the tumour-only pipeline after labelling somatic variants with the three machine learning methods. This procedure was applied to White (134 samples), Black (23 samples), and Asian (16 samples) populations.

### 3.7 Comparison of the TMB with the literature

We compared our strategy to a previously published study [4], using the high-coverage data set. It must be stressed that our ensemble variant calling pipeline integrates two independent variant callers, VarDict and Mutect2, and retains only the variants identified by both tools, in line with best practices in cancer genomics. In contrast, the published study relied on a single proprietary software with undisclosed parameters to construct its gold standard. Consequently, our approach is expected to be, at least in principle, more conservative in terms of variant detection. Consistent with this expectation, our method yielded lower tumour mutational burden (TMB) estimates (Figure 6).

**Fig. 6.**
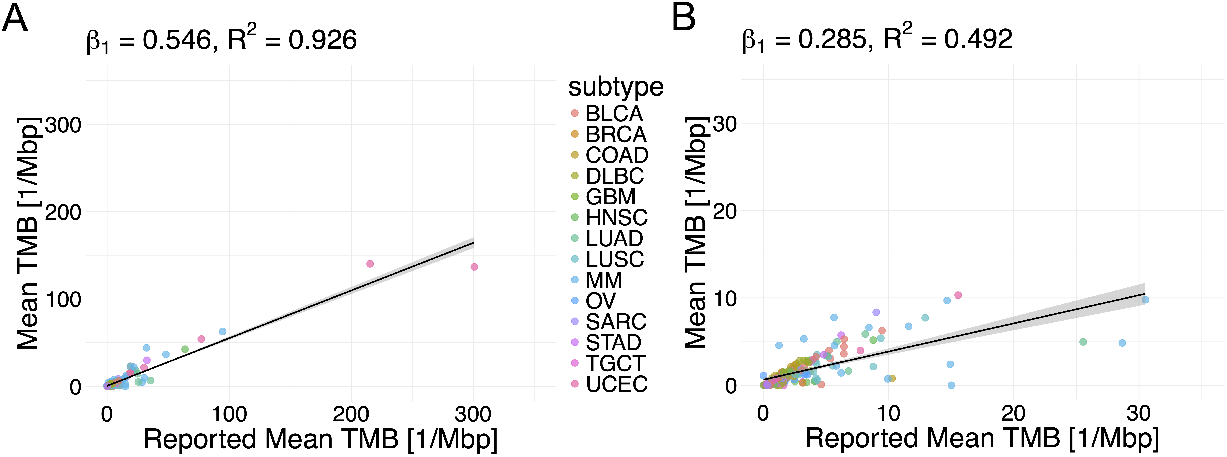
Comparison of our estimate of the TMB (y-axis) with the TMB reported in the literature [4] (x-axis) with the linear regression fit and the corresponding slope *β*_1_ and coefficient of determination (*R*^2^), using both the full data set (A) and after removing outliers (B). The grey area represents the 95% confidence interval. In order to compare our approach with the literature, the TMB was calculated as the mean value of the three models and normalised for a fixed value (41 Mbp) irrespective of the exome capture kit used.

The slope of the linear regression comparing our TMB estimates (y-axis) to those from the literature (x-axis) was consistently below 1, both when considering the full data set (Figure 6A) and when excluding outliers (Figure 6B). Notably, our TMB values also exhibited reduced variability (standard deviation: 13.0 and 27.2, respectively for our estimates and the data reported in the literature), likely reflecting improved robustness to artefacts provided by the combined use of two variant callers.

### 4 Conclusion

Accurately distinguishing somatic mutations from rare germline variants and artefacts in tumour-only whole-exome sequencing (WES) data remains a major challenge in cancer genomics, particularly in the absence of matched-normal samples. This limitation can compromise the reliability of key biomarkers such as the tumour mutational burden, which is increasingly used to guide patient stratification in immunotherapy trials. Existing methods often rely on computationally intensive procedures, including copy-number analysis [32], or suffer from bias introduced by under-representation of certain populations in germline variant databases.

Recent work has shown that machine-learning methods can provide a valuable tool to tackle cancer variant classification [4]. In this work, we developed an open-source, streamlined pipeline for the classification of somatic and rare germline in tumour-only WES data. Our ensemble variant calling method was built on established best practices in cancer variant calling, integrating multiple small variant callers to improve sensitivity and reduce false positives arising from technical artefacts [2]. Importantly, our pipeline avoids computationally demanding steps such as copy-number variation inference [33], allowing for efficient large-scale application without compromising accuracy, and simplifying the engineering of features. Moreover, it does not require the preparation of a panel-of-normal data set, a step which can be computationally intensive. By engineering a concise set of discriminative features, we enabled the robust classification of somatic and rare germline variants through machine learning, making this approach well-suited for both retrospective studies and real-time clinical decision support.

Our workflow demonstrated robustness to ancestry-related bias that commonly affect tumour-only variant calling. By minimizing reliance on population-specific germline databases and instead leveraging a diverse set of orthogonal features, such as variant allele frequency, mutational context, and recurrence in somatic mutation databases, our approach achieved consistent performance across individuals of different genetic backgrounds. This will mitigate the well-documented inflation of TMB estimates in under-represented populations thus supporting more equitable application of genomic biomarkers in precision oncology. Moreover, we demonstrated that our approach generalises across test sets and tumour types more effectively compared to the state-of-the-art, maintaining consistently high accuracy regardless of the data set analysed.

Gradient-boosting trees outperformed deep neural-network models such as TabNet. Consistent with previous reports [34], gradient-boosting methods effectively capture the non-linear relationships present in features derived from DNA sequences or cancer-variant databases, enabling accurate discrimination between somatic and germline states.

In summary, we implemented an open-source pipeline to process tumour-only WES data and used its output to train three machine-learning models tailored to distinguish rare germline variants (with allele frequency *<* 1% in population databases) from somatic variation characterised by variant allele frequency ranging from high to moderate levels. Notably, our method will serve as a reference for future work in the field and is fully available and reproducible. In order to improve the generalisability of our models across diverse data sets, we deliberately excluded certain previously proposed features, such as the total number of variants per sample, to avoid introducing biases specific for a particular data set. We trained our classifiers using a focused set of eight types of features, selected for their robustness and interpretability, instead of the ten types previously reported [4]. Despite this streamlined design, the models achieved high classification accuracy. Notably, high accuracy was maintained even when applied to low-coverage WES data. We observed that tree-based models trained on high-coverage data generalised well to low-coverage test sets, and vice versa, indicating that they are resilient to variations in sequencing depth and suggesting that model performance is driven more by a highly informative and discriminative features than by sequencing depth. Although TabNet baseline accuracy was lower, it also remained remarkably robust to variations in coverage across the training and test data.

As already reported in the literature [4], for somatic mutation calling using DNA sequencing data, we can anticipate that no algorithm will match the performance achieved by using matched-normal samples. Yet, despite the intrinsic limits of interpreting somatic variants against the reference genome rather than its corresponding matched-normal sample, our approach offers a robust solution to estimate the TMB when matched-normal samples are missing.

## Supplementary information

Supplementary Table S1 reports the metadata for all the samples used in the study as well as the TMB values calculated for the high coverage data set. Supplementary Table S2 reports the accuracy metrics for the models training with increasing number of features. Supplementary Table S3 reports the accuracy metrics for the runs with different coverage data and removing important or irrelevant features.

## Acknowledgements

We thank Sakshi Khaiwal and Matteo De Chiara for their valuable feedback.

